# Bioinformatics classification of the MgtE Mg²⁺ channel and de novo protein design for the stabilization of its novel subclass

**DOI:** 10.1101/2025.05.26.656215

**Authors:** Zhixuan Zhao, Kimiho Omae, Wataru Iwasaki, Ziyi Zhang, Fazhi Pan, Eun-Jin Lee, Motoyuki Hattori

## Abstract

MgtE channels play crucial roles in Mg²⁺ homeostasis and are implicated in bacterial survival under antibiotic exposure. Previous structural and biophysical studies have predominantly focused on the *Thermus thermophilus* MgtE, leaving the structural and mechanistic diversity of MgtE family proteins largely unexplored. In this study, using a genome mining approach, we identified diverse MgtE homologs, including a novel subclass termed the “mini-N type,” which lacks the canonical cytoplasmic N and CBS domains but possesses a unique small N-like domain. Despite extensive expression screening, mini-N type homologs could not be stably purified. To address this issue, we designed a series of de novo proteins and determined their crystal structures. A selected de novo protein was fused to a mini-N type MgtE, enabling successful purification and preliminary cryo-EM imaging. Our findings demonstrate that de novo designed protein fusions can serve as powerful tools for stabilizing and purifying otherwise unstable membrane proteins, opening new avenues for the structural and functional studies of otherwise inaccessible membrane proteins.

## Introduction

MgtE represents a widely conserved Mg²⁺ channel in bacteria that plays a critical role in Mg²⁺ homeostasis [1–4] and contributes to bacterial survival under antibiotic exposure by counteracting hyperpolarization [5]. In *Pseudomonas aeruginosa*, MgtE inhibits transcription of the type III secretion system-a critical virulence factor-by promoting the expression of the translation-inhibitory sRNAs RsmY and RmsZ [6, 7]. In humans, MgtE homologues are referred to as SLC41A1 to SLC41A3 and have been implicated in various physiological functions and diseases, including Parkinson’s disease [8–11]. Accordingly, the MgtE/SLC41 family of proteins has attracted considerable interest as a potential therapeutic target.

To date, extensive structural and mechanistic studies have been conducted on MgtE [12–14]. These studies have revealed that MgtE functions as a “Mg²⁺-gated Mg²⁺ channel,” where Mg²⁺ binding to the cytoplasmic domain leads to channel inactivation [3, 15–17]. High-resolution structural analyses of the MgtE transmembrane (TM) domain have elucidated the mechanism of Mg²⁺ selectivity [18, 19], while the cryo-EM structure of MgtE in the absence of Mg²⁺ have provided insights into the gating mechanism [20].

However, these studies have predominantly focused on the *Thermus thermophilus* MgtE (TtMgtE), and structural and biochemical data for MgtE homologues from other species are extremely limited [21, 22]. As a result, the structural and functional diversity of the broadly conserved MgtE family remains largely unexplored. In parallel, the recent explosion in genomic data has made it increasingly feasible to investigate the diversity of specific protein families using genome mining approaches [23, 24].

One major challenge in studying membrane proteins such as MgtE lies in their generally low expression levels and poor stability in detergents [25], which significantly hampers structural and functional analyses. Recently, AI-based de novo protein design technologies, such as ProteinMPNN [26] and RFdiffusion [27], have rapidly advanced [28] and are increasingly expected to facilitate research on membrane proteins [29]. De novo designed proteins typically exhibit high stability, and fusing them to target membrane proteins may enhance expression and stability, potentially enabling the preparation of large quantities of stable membrane protein samples.

In this study, we conducted a bioinformatics analysis of the MgtE family and identified a novel subclass of MgtE proteins with unique domain architectures. Despite extensive expression screening, these proteins could not be stably purified. To overcome this challenge, we performed de novo protein design and successfully determined the crystal structures of several designed proteins. Furthermore, by generating fusion constructs with the de novo designed protein and the novel MgtE subclass, we achieved successful expression and purification. Overall, this study highlights a promising strategy for stabilizing and purifying membrane proteins using a de novo designed protein fusion approach.

## Materials and Methods

### Bioinformatics analysis of MgtE proteins

The prokaryotic proteomes and taxonomic information were downloaded from Genome Taxonomy Database Release 06-RS202 [30]. Only the high-quality (HQ) genomes (completeness ≥95% and contamination ≤5%) based on CheckM [31] were used for the analysis. The genome quality of p Patescibacteria was estimated by using custom marker set provided by CheckM. The hidden Markov model profile of Pfam PF01769 (Divalent cation transporter, MgtE) was used as the query for hmmsearch to find MgtE homologs from the proteomes [32, 33]. The domain architecture of MgtE was determined by InterProScan search [34]. The InterPro IDs IPR006667, IPR046342 and IPR006668 were used as proxies for transmembrane (TM), CBS and N domains of MgtE, respectively.

Data analysis and visualization were performed using R 4.3.2 [35] with the following packages: tidyverse [36], plyranges [37], Biostrings [38] and patchwork [39]. The code for the analysis is provided in the GitHub repository https://github.com/0mae/mgte_short.

### Expression screening of MgtE by FSEC-Nb

MgtE genes and their mutants were synthesized by Azenta (Suzhou, China) and subcloned into a pETNb-nALFA vector [40] containing an N-terminal 8×His tag, an ALFA tag [41], and an HRV 3C protease cleavage site. *Escherichia coli* Rosetta (DE3) cells were transformed with the expression vectors and cultured in 8mL LB medium containing 50 μg/mL kanamycin and 30 μg/mL chloramphenicol at 37°C with the addition of 0.5 mM IPTG when the OD_600_ reached ∼0.6. *E. coli cells* were then cultured at 18°C for 16 hours and harvested by centrifugation. The *E. coli* cell pellets were suspended in 400 μl of buffer TBS (50 mM Tris-HCl pH 8.0, 150 mM NaCl) containing 1 mM phenylmethylsulfonyl fluoride (PMSF) and disrupted by sonication. The cell debris was removed by centrifugation (20,000 × g, 10 min). The supernatant was solubilized with 500 μl of buffer TBS containing 2% (w:v) DDM and 0.5 mM PMSF for 1 hour, followed by ultracentrifugation (200,000 × g, 20 min). 1 μg of EGFP-tagged NbALFA was added to the half amount of the supernatant and incubated for 30 min. After centrifugation (20,000 × g, 10 min), 50 μl of the sample was applied to a Superdex 200 Increase 10/300 GL column or a Superose 6 Increase 10/300 GL column (Cytiva, USA) equilibrated with buffer TBS containing 0.05% (w:v) DDM for FSEC analysis [42, 43]. Fluorescence was monitored using an RF-20Axs fluorescence detector (Shimadzu, Japan) (excitation: 480 nm, emission: 512 nm).

### Expression and purification of MgtE

The large-scale expression of MgtE proteins followed the same method as small-scale expression screening and utilized 6 L of LB medium for expression. After harvesting by centrifugation, the *E. coli* cell pellets were resuspended and disrupted using a microfluidizer in TBS buffer. After centrifugation to remove cell debris, the supernatant was ultracentrifuged at 200,000 × g for 1 hour to isolate the membrane fraction. The membrane was solubilized in S buffer [50 mM Tris (pH 8.0), 150 mM NaCl, 2% n-Dodecyl-β-D-maltoside (DDM), 1 mM PMSF, and 1 mM β-Mercaptoethanol (β-ME)] for 1 hour. After ultracentrifugation at 200,000 × g for 1 hour, the supernatant was loaded onto Ni-NTA agarose beads (QIAGEN, Germany). The beads were washed with 10 column volumes of buffer A (50 mM Tris-HCl [pH 8.0], 150 mM NaCl, 20 mM imidazole, 0.03% DDM, 1 mM β-ME), and proteins were eluted using buffer B (50 mM Tris-HCl [pH 8.0], 150 mM NaCl, 250 mM imidazole, 0.03% DDM, 1 mM β-ME). The eluates were dialyzed against buffer C (20 mM HEPES [pH 7.0], 150 mM NaCl, 0.03% DDM, 1 mM β-ME) for 16 hours at 4°C. The protein was injected onto a Superdex 200 Increase 10/300 GL size-exclusion chromatography column (Cytiva, USA) and eluted using buffer D [20 mM HEPES (pH 7.0), 150 mM NaCl, 0.03% DDM, 0.5 mM Tris(2-carboxyethyl)phosphine (TCEP)].

### De novo protein design

All de novo proteins in this study were designed using the ColabDesign framework (https://github.com/sokrypton/ColabDesign). Briefly, the backbone structures of ZZ1–ZZ3 proteins were generated using RFdiffusion [27] with a specified length of 200 amino acid residues and an unconditional folding mode. The corresponding sequences were then generated using ProteinMPNN [26], and the designed protein sequences were subsequently validated by AlphaFold [44]. The ZZ4-ZZ7 proteins were similarly designed but using the conditional folding mode.

### Expression screening of de novo designed proteins

The DNA sequence encoding each de novo designed protein was synthesized by Azenta (Suzhou, China) and subcloned into a pETNb-cALFA vector [40] containing an HRV 3C protease cleavage site, an ALFA tag, and an 8×His tag at the C-terminus. *E. coli* Rosetta (DE3) cells were transformed with the expression vectors and cultured in 8 mL of LB medium containing 50 μg/mL ampicillin and 30 μg/mL chloramphenicol at 37°C. When the OD_600_ reached ∼0.6, 0.5 mM IPTG was added to induce protein overexpression. The *E. coli* cells were then cultured at 18°C for 16 hours and harvested by centrifugation. The *E. coli* cell pellets were resuspended and disrupted by ultrasonication in 400 μL of TBS buffer [50 mM Tris-HCl (pH 8.0), 150 mM NaCl] supplemented with 1 mM PMSF. After ultracentrifugation (200,000 × g, 20 min), 1 μg of EGFP-tagged NbALFA was added to the supernatant and incubated for 30 min. The mixture was then centrifuged at 20,000 × g for 10 min, and 50 μL of the sample was applied to a Superdex 200 Increase 10/300 GL column (Cytiva, USA) equilibrated with TBS buffer for the FSEC assay.

### Expression and purification of de novo designed proteins

The large-scale expression of de novo designed proteins followed the same method as small-scale expression screening, utilizing 6 L of LB medium for expression. After harvesting by centrifugation, the *E. coli* cell pellets were resuspended in TBS buffer. Following centrifugation, the supernatant was loaded onto Ni-NTA agarose beads and washed with 10 column volumes of buffer E (50 mM Tris-HCl, pH 8.0, 150 mM NaCl, 20 mM imidazole). Proteins were then eluted using buffer F (50 mM Tris-HCl, pH 8.0, 150 mM NaCl, 250 mM imidazole). After elution, the ALFA tag and 8×His tag were cleaved by HRV 3C protease while the eluates were dialyzed against buffer G (20 mM HEPES, pH 7.0, 150 mM NaCl) for 16 h at 4°C. The sample was then reapplied to Ni- NTA beads, and the flow-through fractions were injected onto either a Superdex 200 Increase 10/300 GL or a Superdex 75 Increase 10/300 GL size-exclusion chromatography column (Cytiva, USA) in buffer G. The peak fractions were concentrated to 10 mg/mL for crystallization.

### Crystallization

One microliter of ZZ1 protein was mixed with 1 μl of crystallization reservoir buffer H [0.05 M zinc acetate, 0.05 M MES, pH 6.0, 7.6% PEG 8000], while 1 μl of ZZ4 protein was mixed with 1 μl of crystallization reservoir buffer I [0.2 M MgCl₂, 0.1 M KCl, 0.025 M sodium citrate, pH 4.0, 24.5% PEG 400] for crystallization using the vapor diffusion method at 18°C. Crystals typically appeared within one week and were subsequently harvested in reservoir solutions supplemented with a final concentration of 30% glycerol for ZZ1 or 38% PEG 400 for ZZ4, respectively before flash-freezing in liquid nitrogen for X-ray diffraction experiments.

### X-ray data collection and structure determination

X-ray diffraction experiments for ZZ1 and ZZ4 were performed at beamline BL02U1 of the Shanghai Synchrotron Radiation Facility (SSRF) [45]. Data processing was carried out using XDS [46]. The initial phase was determined by molecular replacement using Phaser [47], utilizing AlphaFold [44] models as templates. Manual model building and iterative refinement were performed with Coot [48] and PHENIX [49], respectively. Structural figures were prepared using PyMOL (https://pymol.org).

### Construct design of ZZ1-fused VhMgtE

The ZZ1-fused VhMgtE harbors a 3- or 4-residue truncation at their C-terminus and are fused via one or two EK2R1 (AEEEKRK) helices [50]. The construct was validated by AlphaFold [44] before wet experiments.

### Cryo-EM of ZZ1-fused MgtE

The purified ZZ1-fused MgtE protein was mixed with Amphipol A8-35 (Anatrace, USA) at a mass ratio of 1:10 and incubated for 16 hours. Bio-Beads SM-2 (Bio-Rad, USA) were then added to the mixture and incubated at 4°C for 4 hours to remove detergents. The complex was further purified by size-exclusion chromatography using a Superdex 200 10/300 column in buffer J (20 mM HEPES pH 7.0, 150 mM NaCl, 0.5mM TCEP). The peak fractions were concentrated to 2.0 mg/mL. Freshly prepared samples were incubated with 2 mM MgCl₂ on ice for 30 minutes before vitrification. Cryo-grid preparation was performed at 4°C and 100% humidity using a Vitrobot Mark IV (Thermo Fisher, USA). A 2.5 µL sample was applied to each freshly glow-discharged Au 300-mesh grid (Quantifoil, Germany, R1.2/1.3). The grids were then plunge-frozen in liquid ethane. Cryo-grids were screened using a 200 kV Glacios 2 microscope equipped with a Falcon 4i camera (Thermo Fisher, USA).

### Data and code availability

The atomic coordinates of the de novo designed protein structures were deposited in the Protein Data Bank under accession code ID: 9LNI (ZZ1) and 9LX4 (ZZ2). All relevant data are included in the paper, its supplemental information or deposited in Mendeley Data (https://data.mendeley.com/datasets/kdns5hnrmw/1) and GitHub (https://github.com/0mae/mgte_short).

## Results and Discussion

### Bioinformatics analysis of MgtE proteins identified the mini-N type of MgtE

We conducted an extensive search against prokaryotic genome data sets containing 27,742 genomes to identify genes encoding the TM domain of MgtE, which revealed a variety of domain architectures (Figure 1). The canonical type of MgtE (e.g., *Thermus thermophilus* MgtE) contains all three domains: the N domain, the CBS domain, and the TM domain (N-CBS-TM; Figure 1A). However, some MgtE proteins lack the N domain (CBS-TM), while others lack both the N and CBS domains, retaining only the TM domain (TM; Figure 1B). These proteins are not negligible in number (∼2,600), and they were conserved in both bacteria and archaea (Table 1). Notably, some proteins lacking both the N and CBS domains possess tandem-repeat TM domains (TM-TM), similar to the eukaryotic MgtE homolog, the SLC41 family (Figure 1C).

**Fig. 1.**
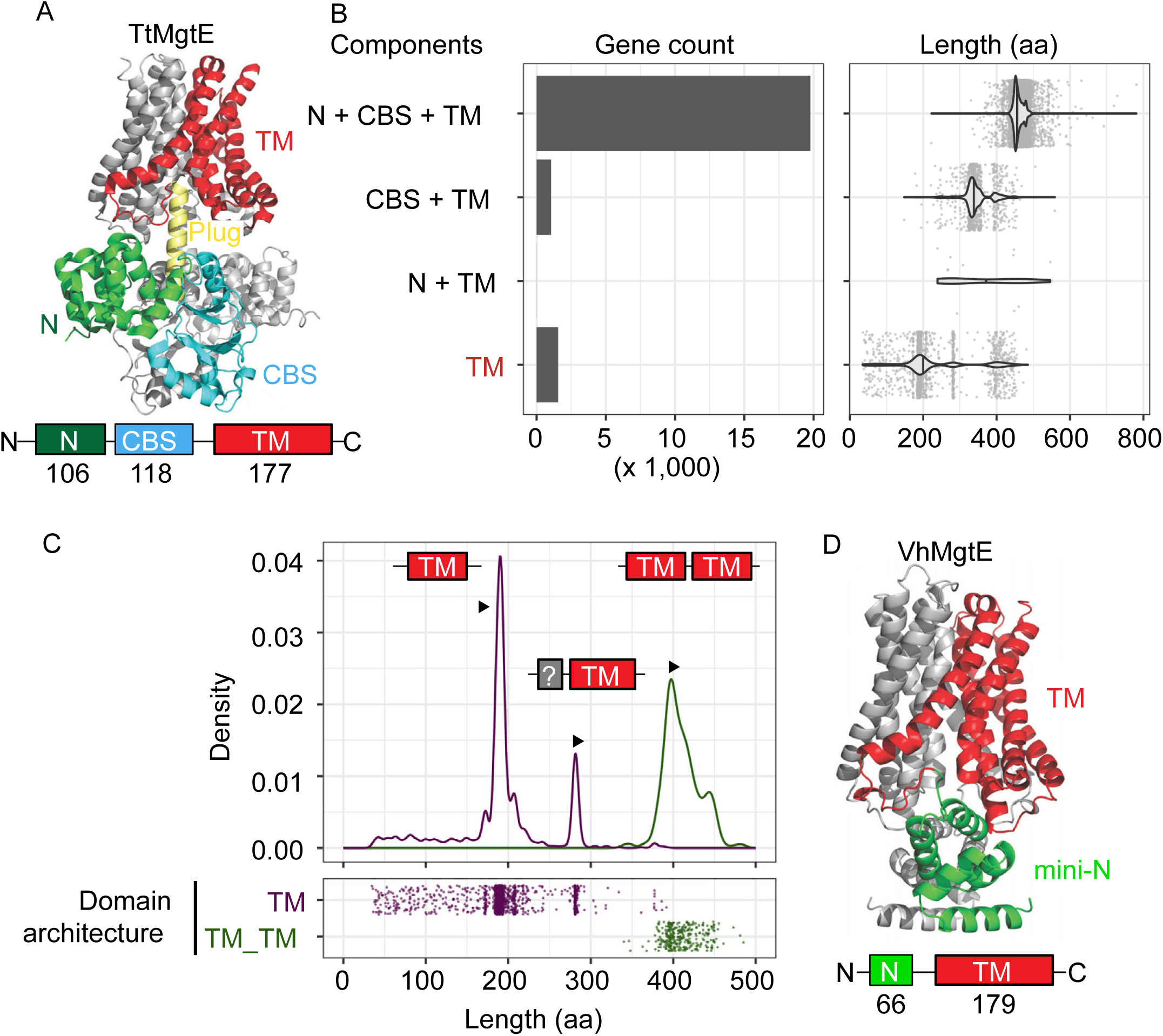
Bioinformatics analysis of MgtE proteins identified various types of MgtE. (A) The canonical MgtE structure. The structure of *Thermus thermophilus* MgtE (PDB ID: 2ZY9). (B) The number and length of MgtE proteins identified by bioinformatics analysis. MgtE proteins are classified based on the domain components. (C) The length distribution of MgtE gene that lacks both the N and CBS domains. MgtE proteins are classified based on the number of TM domains. The novel MgtE protein is indicated by the red arrow. (D) The structure of the novel “mini-N” type MgtE. The structure of *Vibrio harveyi* MgtE (UniProt ID: A0A0D0IA63) predicted by AlphaFold2.

**Table 1.** The number of MgtE homologs.

Intriguingly, we also identified a novel type of MgtE gene that lacks both the N and CBS domains but contains an additional domain structurally similar to, yet smaller than, the canonical N domain (Figure 1C and S1). We designated this class of MgtE as the “mini-N” type (Figure 1D). The discovery of the mini-N-type MgtE was surprising, as in the canonical MgtE, the CBS domain is known to be essential for channel regulation, while the N domain itself does not appear to have a direct role in gating regulation[3].

### Expression screening of the mini-N type MgtE proteins

To structurally and functionally characterize these various types of MgtE proteins, we initially selected mini-N-type MgtE proteins for expression screening before undertaking large-scale expression and purification. We expressed 12 genes encoding mini-N-type MgtE proteins in E. coli at a small scale and evaluated their expression profiles using fluorescence-detection size-exclusion chromatography (FSEC) [42, 51], specifically through nanobody (Nb)-based FSEC (FSEC-Nb)[40] (Figure 2). In the FSEC-Nb method, targeted membrane proteins are fused to an ALFA peptide tag [41], and solubilized whole-cell extracts were mixed with anti-ALFA tag Nb (NbALFA) fused to GFP for subsequent FSEC analysis to detect the complex of the target membrane protein and the Anti-ALFA tag NB fused to GFP. FSEC-Nb screening of mini-N-type MgtE homologs showed that multiple homologs (Nos. 5–9 in Figure 2) exhibited sharp, high-intensity peaks, indicating the high level of expression and monodispersity, so that they might be suitable for further large-scale expression and purification. However, in further expression and purification trials, these proteins showed a tendency to aggregate or precipitate during purification.

**Fig. 2.**
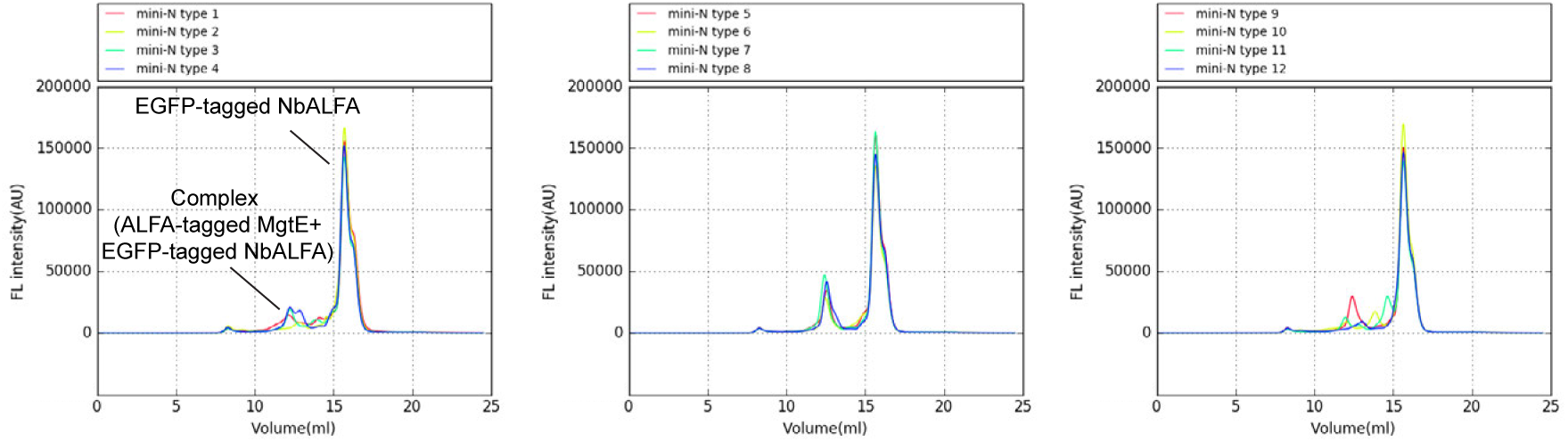
Expression screening of mini-N type MgtE proteins using FSEC-Nb. FSEC traces of unpurified ALFA-tagged mini-N type MgtE proteins mixed with EGFP-tagged NbALFA on a Superdex 200 Increase 10/300 GL column, detected via EGFP fluorescence. Expression of mini-N type MgtE proteins from the following species was screened: 1: *Endozoicomonas elysicola* (WP_081622142.1), 2: BD1-7 clade bacterium (CAA0079619.1), 3: *Endozoicomonas sp*. OPT23 (WP_228550828.1), 4: *Endozoicomonas numazuensis* (WP_034843082.1), 5: *Photobacterium frigidiphilum* (WP_107243853.1), 6: *Enterovibrio baiacu* (WP_129495674.1), 7: *Vibrio thalassae* (WP_096991926.1), 8: *Vibrio* (Multispecies) (WP_146490653.1), 9: *Vibrio harveyi* BSW5 (A0A0D0IA63), 10: *Gloeocapsa sp.* PCC 7428 (WP_015189061.1), 11: *Nodularia sp.* NIES-3585 (WP_089089827.1), 12: *Pandoraea terrae* (WP_150700210.1).

### Design of de novo proteins

To overcome these challenges in the purification of mini-N-type MgtE homologs, we designed a series of de novo proteins (Figure 3 and Table S1) to serve as potential fusion partners. Protein fusion strategies have been widely employed for membrane protein stabilization and structural studies [52, 53], as fusion partners can not only enhance the stability of membrane proteins but also facilitate crystallization and particle alignment during cryo-EM data processing. Furthermore, de novo designed proteins generally exhibit extremely high stability [27, 28], potentially making them particularly suitable as fusion partners for membrane proteins. Using RFdiffusion [27], we designed seven de novo proteins: three (ZZ1–ZZ3) using unconditional folding mode with a fixed length of 200 amino acid residues, and four (ZZ4–ZZ7) using conditional folding mode with lengths close to 200 residues. Validation through AlphaFold2 prediction yielded high pLDDT and pTM scores, with small root mean square deviation (RMSD) values ranging from approximately 0.4 to 0.6 Å (Table S1). In the unconditional folding mode, we generated all-α (ZZ1), α/β (ZZ2), and α+β (ZZ3) structures (Figure 3A-C). In the conditional folding mode, besides an all-α structure (ZZ5), we aimed to design β-barrel architectures containing internal α-helix elements (ZZ4, ZZ6, and ZZ7), reminiscent of the GFP fold [54]. Intriguingly, despite their relatively simple topologies, none of these designed proteins exhibited significant overall structural similarity to proteins in the PDB100 database as analyzed by FoldSeek [55] (minimum structural alignment overlap of 90% and an E-value cutoff of 0.01 as a threshold). These observations support the idea that the structural space accessible by de novo designed proteins is far broader than the space currently represented by naturally occurring protein structures [56].

**Fig. 3.**
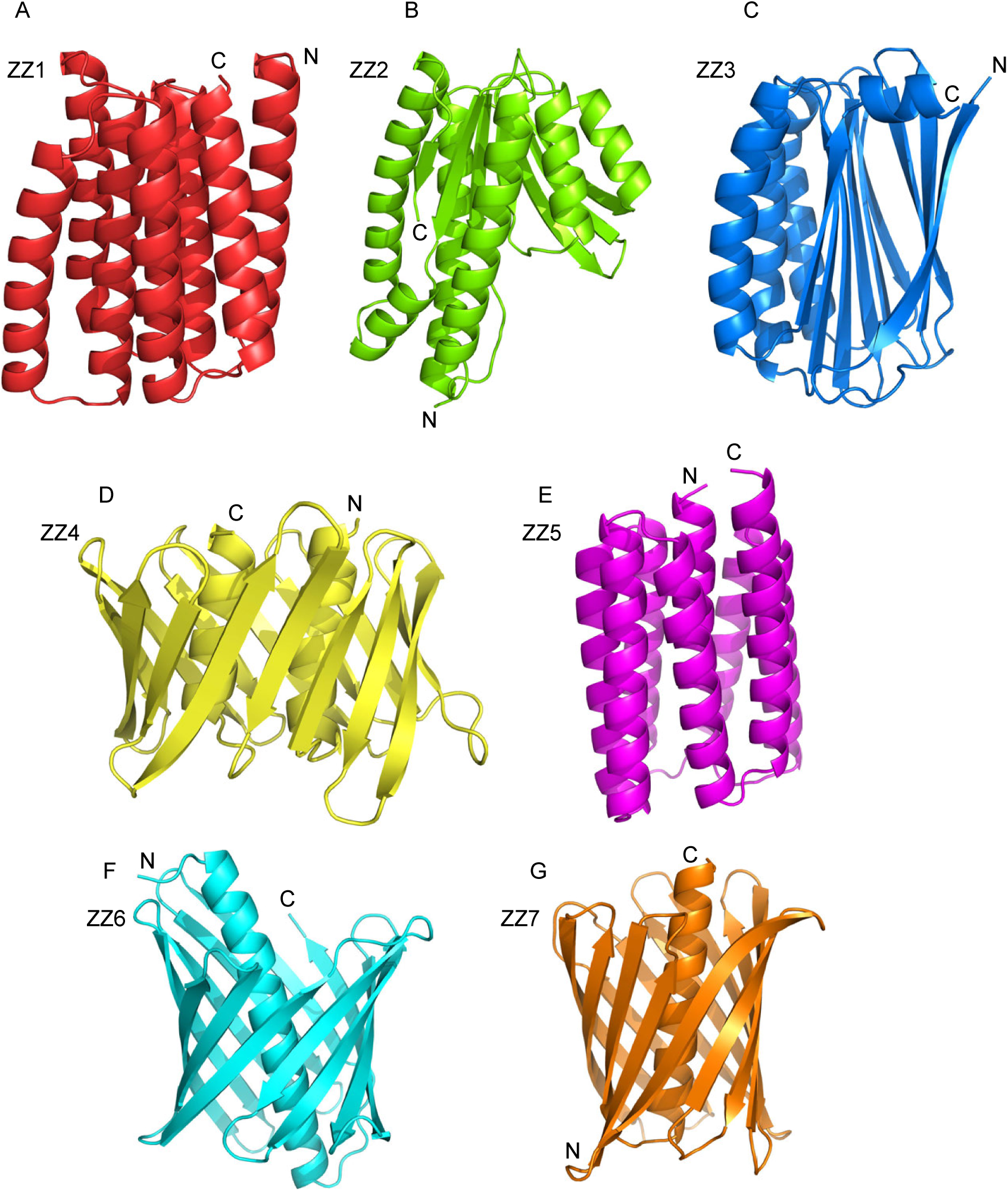
De novo protein design using RFdiffusion. (A–G) Cartoon representations of de novo designed protein structures predicted by AlphaFold2.

### Experimental verification of de novo designed proteins

The designed proteins were screened for expression in *E. coli* using the FSEC-Nb method [40] prior to their structural analyses (Figure 4A). Although FSEC techniques are primarily utilized for screening membrane protein expression, they can also be effectively applied to soluble proteins [51]. All of seven designed proteins showed high and sharp peaks corresponding to the complex of the ALFA-tagged these proteins and NbALFA fused to GFP in FSEC (Figure 4A), and were subsequently purified for crystallization trials. Notably, the elution profiles of these proteins corresponded to monomeric molecular weights, as anticipated from our design. Crystals of ZZ1 and ZZ4 diffracted to resolutions of 1.61 and 2.03 Å, respectively, and their structures were successfully determined by molecular replacement using AlphaFold2 models as templates (Figure 4B and Table S2). Structural superposition revealed RMSD values of 0.65 Å and 0.83 Å for the Cα atoms of ZZ1 and ZZ4, respectively, between the AlphaFold-predicted models and crystal structures, demonstrating the high accuracy of our designs (Figure 4B).

**Fig. 4.**
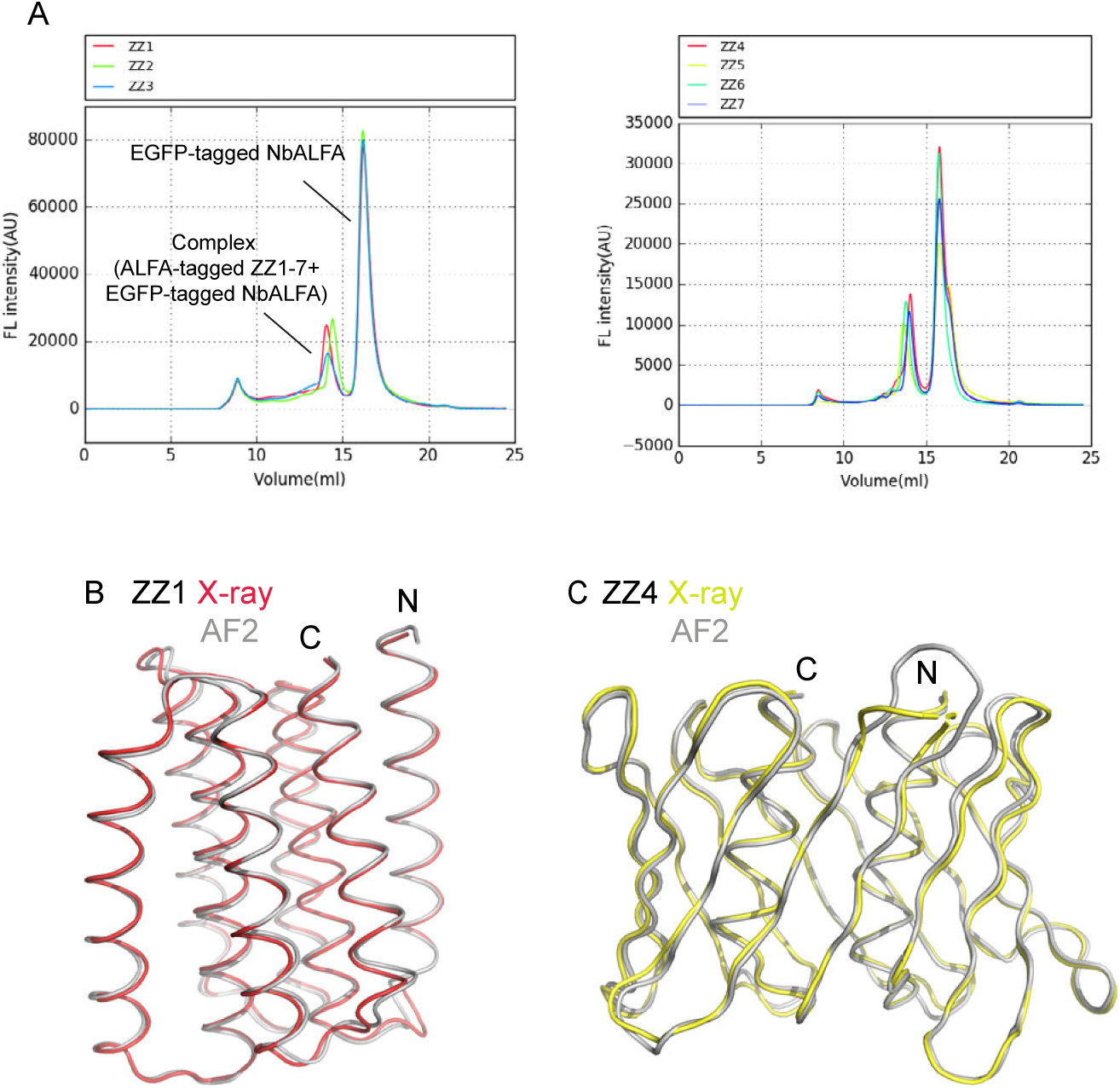
Expression screening and structural analysis of de novo designed proteins. (A) FSEC traces of unpurified de novo designed proteins ZZ1–ZZ7 mixed with EGFP-tagged NbALFA on a Superdex 200 Increase 10/300 GL column, detected via EGFP fluorescence. (B–C) Crystal structures of de novo designed proteins ZZ1 (B) and ZZ4 (C). The AlphaFold2-predicted structures are superimposed and shown in gray.

### Fusion of the de novo designed protein enabled the mini-N type MgtE purification

To fuse our de novo designed proteins to the mini-N type MgtE, we selected ZZ1 based on its α-helix bundle architecture, which is highly compatible with fusion to membrane proteins in various configurations, as well as its demonstrated stability evidenced by successful X-ray crystallography (Figure 4). For the mini-N type MgtE, we chose the homolog from Vibrio harveyi (VhMgtE) due to its favorable peak profile observed in FSEC-Nb analysis (No. 9 in Figure 2).

To fuse ZZ1 to VhMgtE, we employed an EK2R1 (AEEEKRK) helical linker (Figure 5A), known for its high stability [57] and utility in membrane protein structural determination by fusion strategies [58]. We designed three constructs, fusing ZZ1 to the C-terminus of VhMgtE: Construct 1 (3-residue truncation at the C-terminus of VhMgtE with one EK2R1 linker), Construct 2 (3-residue truncation at the C-terminus of VhMgtE with two EK2R1 linkers), and Construct 3 (4-residue truncation at the C-terminus of VhMgtE with two EK2R1 linkers), as validated by AlphaFold2 structure prediction (Figure S1). These constructs were then screened for expression using FSEC-Nb (Figure 5B).

**Fig. 5.**
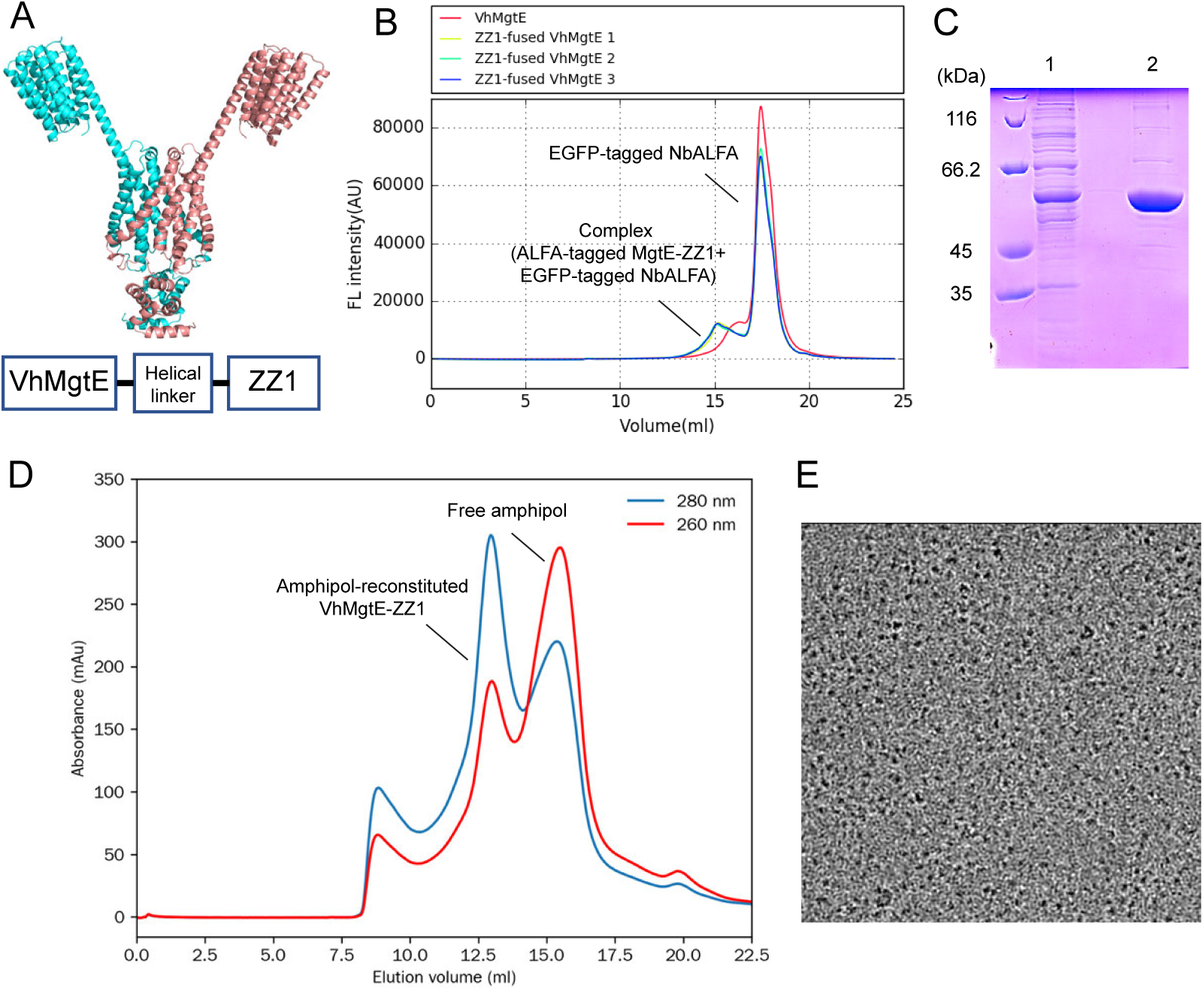
Purification and preliminary cryo-EM of ZZ1-fused VhMgtE. (A) AlphaFold-predicted structure (Construct 3) and domain composition of the ZZ1-fused VhMgtE construct. (B) FSEC traces of unpurified ZZ1-fused VhMgtE constructs mixed with EGFP-tagged NbALFA on a Superose 6 Increase 10/300 GL column, detected via EGFP fluorescence. (C) SDS-PAGE of purified ZZ1-fused VhMgtE construct 3. Lane 1: Flowthrough from Ni-NTA affinity chromatography. Lane 2: Elution fraction. (D) Size-exclusion chromatography of ZZ1-fused VhMgtE construct 3 after amphipol reconstitution, detected via UV absorbance. (E) Cryo-EM image of ZZ1-fused VhMgtE construct 3.

All three constructs exhibited expression levels similar to that of the wild type (Figure 5B). We initially selected Construct 3 for purification. Construct 3 was successfully purified (Figure 5C), reconstituted into amphipol A8-35, and showed a sharp, monodisperse peak in SEC analysis (Figure 5D). Furthermore, preliminary cryo-EM imaging of amphipol-reconstituted ZZ1-fused VhMgtE, using a 200 kV Glacios 2 microscope, showed monodisperse particles (Figure 5E).

Taken together, our results showed that the fusion strategy utilizing de novo designed proteins effectively enabled the purification of this novel subclass of MgtE protein.

### Concluding remarks

Our bioinformatics pipeline identified a novel “mini-N” subclass of MgtE Mg²⁺ channels with atypical domain architectures, suggesting previously unrecognized structural diversity within the MgtE family. While we encountered challenges in expression and purification of the mini-N type MgtE, we tackled these technical issues using AI-driven de novo protein design and protein engineering. We designed stable de novo proteins using state-of-the-art deep learning tools and determined the crystal structures of selected constructs, validating the design approach. Fusion of one such de novo protein (ZZ1) to a mini-N type MgtE homolog significantly enabled purification suitable for structural studies. Preliminary cryo-EM analysis further confirmed the structural homogeneity of the fusion construct. Overall, our work highlights the utility of AI-assisted de novo protein design in overcoming key bottlenecks in membrane protein research. The fusion strategy we developed may be broadly applicable to other challenging targets, paving the way for deeper insights into membrane protein structure and function.

## Acknowledgments

We thank the staff from BL02U1 at Shanghai Synchrotron Radiation Facility (SSRF) for assistance during X-ray data collection. The diffraction experiments were performed at SSRF BL02U1 (Proposal No. 2020-SSRF-PT-014702). We appreciate Ruiliang Jia in Hattori lab for the technical support regarding the FSEC screening of de novo designed proteins. We also appreciate the kind support of Dr. Wataru Iwasaki as a former senior mentor to Kimiho Omae. Computation time was provided by the SuperComputer System, Institute for Chemical Research, Kyoto University.

---

**Fig. S1.**
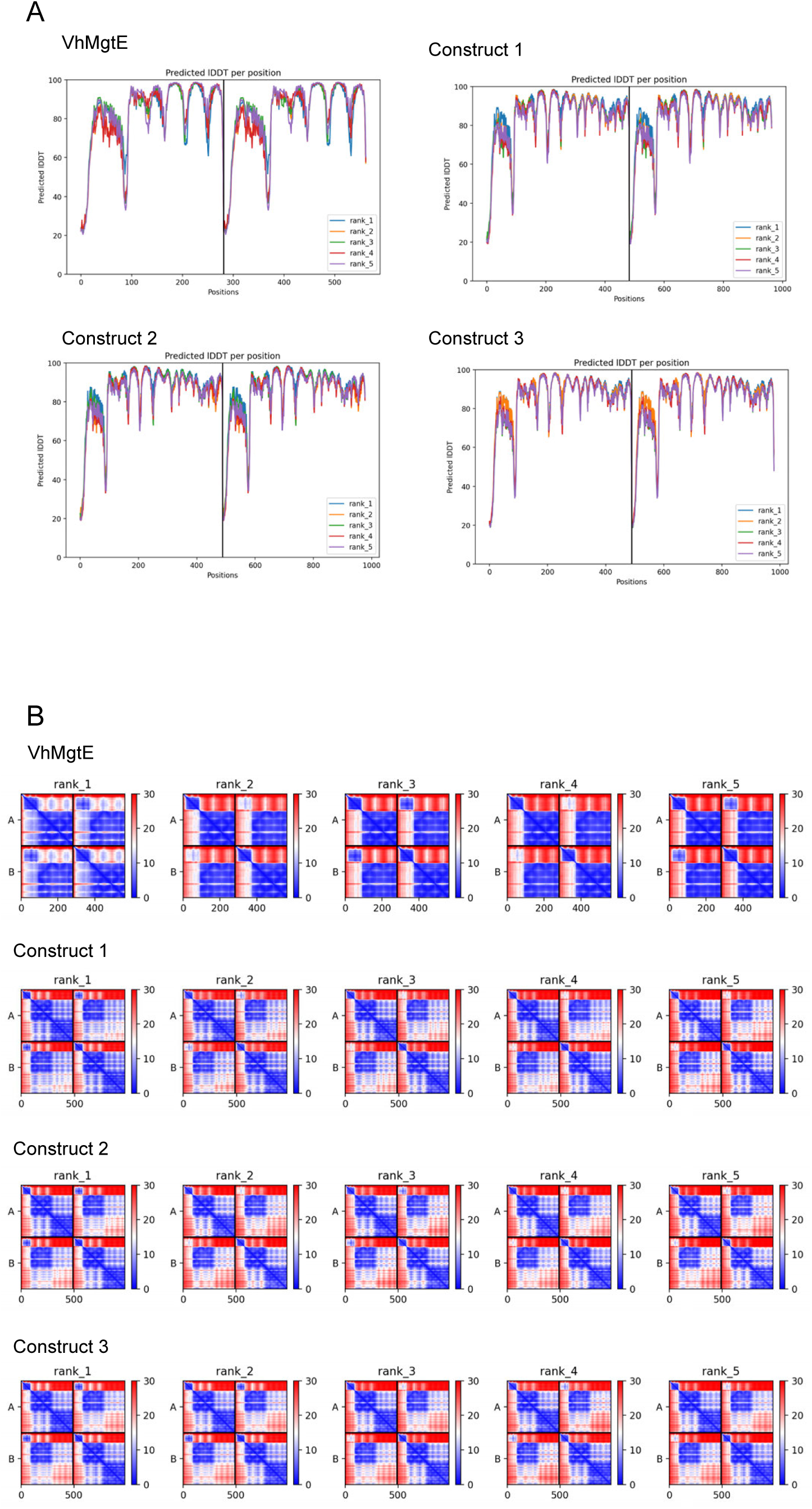
AlphaFold2 statistics of VhMgtE constructs. (A) pLDDT scores for each amino acid position. (B) PAE maps.

**Table S1. Sequence and AlphaFold2 statistics of de novo designed proteins**

**Table S2. X-ray data collection and refinement statistics**

