## Supplementary figures and images for "Bioinformatics classification of the MgtE Mg²⁺ channel and de novo protein design for the stabilization of its novel subclass"

### Figure S1

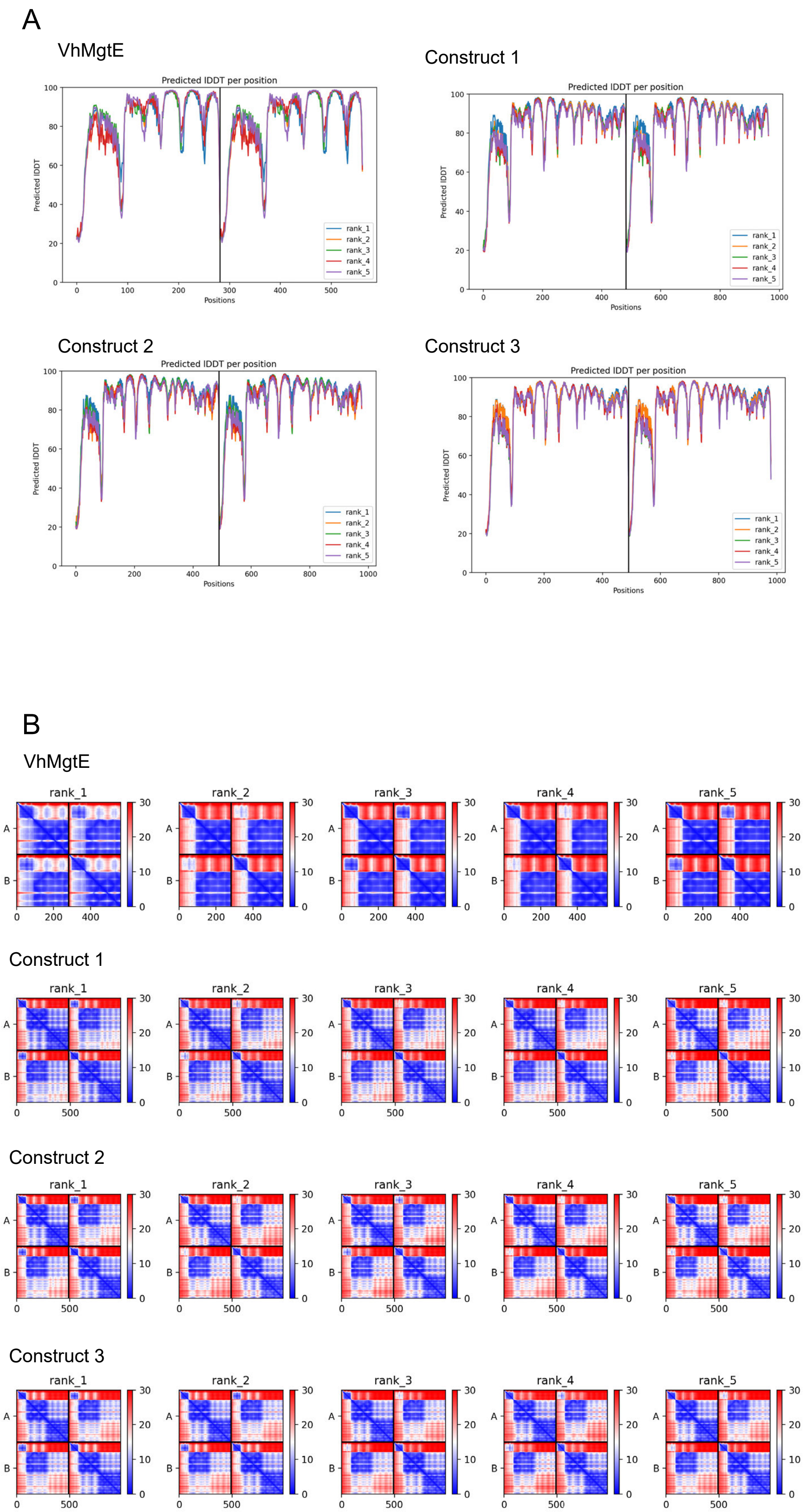
