## Supplementary material for "Bioinformatics classification of the MgtE Mg²⁺ channel and de novo protein design for the stabilization of its novel subclass": Table S2

**Table S2 X-ray data collection and refinement statistics**

|  | ZZ1 | ZZ4 |
| --- | --- | --- |
| **PDB ID** | 9LNI | 9LX4 |
| **Data collection** |  |  |
| Wavelength (Å) | 0.9792 | 0.9792 |
| Space group | *P*2_1_ | *P*2_1_ |
| Cell dimensions |  |  |
| *a*, *b*, *c* (Å) | 35.3, 67.4, 43.7 | 57.4, 58.3, 67.3 |
| *α*, *β*, *γ* (°) | 90.0, 108.5, 90.0 | 90.0, 112.1, 90.0 |
| Resolution (Å)^*^ | 41.48 – 1.61 (1.70 – 1.61) | 62.38 – 2.03 (2.14 – 2.03) |
| *R*_merge_^*^ | 0.064 (0.506) | 0.112 (0.998) |
| *I*/σ*I*^*^ | 8.3 (2.5) | 6.4 ( 1.6) |
| Completeness (%)^*^ | 97.3 (95.5) | 100.0 (99.8) |
| Redundancy^*^ | 4.4 (4.4) | 4.1 (3.2) |
| CC_1/2_ (%)^*^ | 99.7 (86.7) | 99.3 (50.1) |
| **Refinement** |  |  |
| Resolution (Å) | 1.61 | 2.03 |
| No. reflections | 24303 | 26840 |
| *R*_work/_ *R*_free_ | 0.209/0.243 | 0.223/ 0.290 |
| No. atoms |  |  |
| Protein | 1739 | 2587 |
| Other | 7 | 15 |
| Water | 163 | 106 |
| B-factors |  |  |
| Protein | 28.32 | 41.99 |
| Other | 42.89 | 69.47 |
| Water | 39.13 | 49.85 |
| R.m.s deviations |  |  |
| Bond lengths (Å) | 0.008 | 0.008 |
| Bond angles (°) | 1.102 | 1.088 |
| Ramachandran plot |  |  |
| Favoured (%) | 100.0 | 100.0 |
| Allowed (%) | 0.0 | 0.4 |
| Outliers (%) | 0.0 | 0.0 |

*Highest resolution shell is shown in parenthesis.
